# Multiplex high resolution melting PCR for simultaneous genotyping of pyrethroid-resistance associated mutations in *Aedes aegypti.* First report on kdr mutations in wild populations from Argentina

**DOI:** 10.1101/2022.08.26.505371

**Authors:** Alberto N. Barrera Illanes, María Victoria Micieli, Marina Ibáñez Shimabukuro, Soledad Santini, Ademir J. Martins, Sheila Ons

## Abstract

**Background:** *Aedes aegypti* is an urban mosquito vector of Dengue and other arboviruses. During epidemic periods, pyrethroid insecticides are used for the control of adult mosquitoes; the worldwide distributed resistance to these insecticides is a cause of failures in vector control campaigns. The primary target of pyrethroids is the voltage-gated sodium channel; point mutations on this channel, called kdr mutations, are associated with pyrethroid resistance. Two *kdr* mutations, called V1016I and F1534C, augmented in frequency in natural populations of *Ae. aegypti* from the Americas in the last decade. The diagnostic of *kdr* polymorphisms allows an early detection of insecticide resistance spreading, which is critical for timely decisions on vector management. Given the relevance of resistance management, high-throughput methods for *kdr* genotyping are invaluable tools for resistance monitoring programs. These methods should also be cost-effective, to allow regional-scale surveys. Despite the extended presence of *Ae. aegypti* and the incidence of dengue in Argentina, the presence, abundance and distribution of kdr mutations were not reported in this country up to date.

**Methodology and findings:** We report a multiplex high-throughput assay based in High Resolution Melting PCR for the simultaneous genotyping of 1016 and 1534 sites in voltage-gated sodium channel gene. We used this method for the study of individual mosquito samples collected in localities which received different selection pressure with pyrethroids. Compared to other genotyping methods, multiplex High Resolution Melting was high-throughput, cost-efficient, sensitive and specific. We demonstrate for the first time the presence of *kdr* mutations in Argentina in regions under different selection pressure with pyrethroids.

**Conclusions and Significance:** We have developed a high-throughput method for the genotyping of alleles associated with pyrethroid resistance in *Ae. aegypti* from the American continent. The method developed here is comparable in its sensitivity and reliability with other genotyping methods, but reduces costs and running time. It could be incorporated in control campaigns for control the presence and spreading of resistance-associated alleles. We report here for the first time the presence of kdr mutations in distant populations from Argentina, with different epidemiological situations and different history of mosquito control efforts.

**Authors summary:** *Aedes aegypti* is a mosquito vector of viruses such as dengue, causing millions of infections yearly worldwide. Emergence and distribution of insecticide resistance in this mosquito is a challenge for control campaigns. In this context, the implementation of resistance management strategies became a requisite for successful and sustainable mosquito management. To achieve this objective, it is important to count with early genetic markers of resistance, such as *kdr* mutations, which are associated with pyrethroid resistance phenotype. Kdr markers can be detected by molecular genotyping diagnostic at early stages in the emergence of resistance; this information should be considered in rational design of control campaigns. Here we present a high-throughput cost-effective method to detect kdr mutations in *Ae. aegypti*. Also, we demonstrate for the first time the presence of kdr mutations in Argentina.

## Introduction

Around 390 million people in the world are infected by dengue virus every year. In the Americas, the number of reported cases has been multiplied by 10, comparing the 1.5 million cases in the ‘80s to more than 16 million in the last decade [1]. Dengue, zika, chikungunya and the re-emerging yellow fever viruses are transmitted by *Aedes aegypti* female mosquitoes; this species is considered one of the most successful invasive species worldwide [2]. Urbanization, human population growth and climate change are factors that promote the expansion and abundance of *Ae. aegypti* [3]. This mosquito was reintroduced into Argentina in 1986 and is now established in all the northern and central provinces, its southern boundary being the provinces of Buenos Aires [4], Neuquén [5] and Río Negro [6]. Since then, three major and wide distributed dengue epidemics have occurred in Argentina (years 2009, 2016, and 2020), representing a considerable public health concern. The recent outbreak occurred during 2019-2020 period caused more than 67.000 cases https://www.argentina.gob.ar/salud/epidemiologia/boletines [Accessed July 2020 and April 2022].

The control of *Ae. aegypti* expansion consists in the environmental management by the elimination of domestic and peri-domestic larvae breeding sites and the use of larvicides such as insect grow regulators, organophosphate neurotoxics and the biolarvicide *Bacilus thuringiensis israelensis* [7]. The use of adulticides is recommended to be restricted to arbovirus epidemic periods [7]; for this, the neurotoxic pyrethroids are the preferred compounds given their favorable toxicological properties. Even though the organophosphate malathion is used in cases of high pyrethroid resistance in other countries, its domi-sanitary application in Argentina is banned [8].

The target site of pyrethroid insecticides is the voltage-gated sodium channel (Na_v_), a membrane protein present in excitable cells. Given their rapid action, the effect of pyrethroids is known as *knockdown*; the single nucleotide polymorphisms in the sodium channel gene that confer resistance to pyrethroids are known as knockdown resistance (*kdr*) mutations [9]. In the region of the Americas, three *kdr* mutations related to the loss of pyrethroid susceptibility have been identified in *Ae. aegypti*: Ile to Met in position 1011 (I1011M), Val to Ile in position 1016 (V1016I) and Phe to Cys in position 1534 (F1534C) [10]. The frequency of I1011M seems to decrease in the last 20 years, and its role in the pyrethroid resistance is not clear for natural populations [11]. Conversely, V1016I and F1534C mutations augmented in frequency in natural populations in the Americas in the last decade [11–13]. Considering 1016 and 1534 positions, three alleles of *Ae. aegypti* Na_v_ gene (*aedaenav*) are widely distributed in South and North America [12,14,15]: 1016 V + 1534 F (Susceptible; Na_v_S); 1016 V + 1534^kdr^C (Resistant 1; Na_v_R1); 1016^kdr^ I +1534^kdr^ C (the double mutant Resistant 2; Na_v_R2). The Na_v_R3 allele (1016^kdr^ I +1534 F) was detected with a very low frequency (≤ 0.1% of the samples) in Brazil [16], México [12]and Florida [14]. Conversely, the Na_v_R1 and Na_v_R2 genotypes are predominant in all the regions studied in the American continent.

The role of kdr mutations in pyrethroid resistance in *Ae. aegypti* has been well established. In the last 10 years, kdr alleles in natural populations propagated in parallel to increased levels of resistance to pyrethroids [14–16]. More recently, homozygous Na_v_R1R1, Na_v_R2R2 and the heterozygous Na_v_R1R2 laboratory lines of *Ae. aegypti* were compared for their sensitivity to deltamethrin with a Na_v_SS susceptible line, confirming that the three kdr genotypes conferred deltamethrin resistance to the insects. From them, the Na_v_R2R2 genotype conferred the higher levels of resistance [17]. Electrophysiological records in *Xenopus laevis* oocytes showed reduced affinity to type I (permethrin) but not type II (deltamethrin) pyrethroids in oocytes expressing F1534C^kdr^ sodium channel (Na_v_R1 allele) compared to those expressing the Na_v_SS [18]. In parallel, channels expressing the Na_v_R3 allele did not present reduced sensitivity to pyrethroids when expressed in *X. laevis* oocytes. Moreover, when Na_v_R2 allele was expressed in *X. laevis* oocytes, a higher resistance to both type I and type II pyrethroid was observed, when compared to the NavR1 allele [19].

Resistance monitoring is critical for the success of vector control campaigns, and must be considered for operational decision making. The diagnostic of kdr polymorphisms allows an early detection of insecticide resistance spreading, given that it can detect a carrying individual before the mutation is fixed in the population by the selection pressure exerted by the insecticide. For this, studies on kdr distribution and abundance have been performed in American countries such as Venezuela [20], Mexico [12], USA [14] and Brazil [15,16,21]. Despite the extended presence of *Ae. aegypti* and the incidence of dengue in Argentina the existence, abundance and distribution of kdr mutations were not reported in this country up to date.

Given the relevance of resistance management, cost-effective and high-throughput methods for *kdr* genotyping are invaluable tools for resistance monitoring programs. For *Ae. aegypti*, the methods implemented to date are based either on allele-specific PCR [22], or in the use of TaqMan probes [15]. While the economic running cost of allele specific PCR is minor than TaqMan [23], it could be less reproducible by different laboratories, given the requirement of a careful setting of PCR conditions to avoid nonspecific amplifications. On the other hand, High Resolution Melting (HRM) is a reproducible probe-free high throughput method for genotyping. It uses well-calibrated equipment and a third generation fluorescent dsDNA dye [24]. HRM revealed economical and practical advantages with respect to other methods for the detection of *kdr* in *Anopheles gambiae* [23]. Even though single-site HRM was used for genotyping *kdr* mutations in Asian populations of *Ae. aegypti* [25], a development of a multiplex HRM-based assay (mHRM) for *kdr* genotyping in this mosquito species has not been reported to date, up to our knowledge. Here, we report a mHRM assay for the simultaneous genotyping of 1016 and 1534 sites in the *aenav* gene. Compared to other genotyping methods, mHRM was high-throughput, cost-efficient, sensitive and specific. We used this method for the study of individual mosquito samples collected in Argentinean localities which received different selection pressure with pyrethroids, demonstrating for the first time the presence of *kdr* mutations in Argentina in regions under different selection pressure with pyrethroids.

## Methods

### Mosquito sampling and DNA extraction

Mosquito sampling was performed in 2018 and 2019 in the provinces of Salta (Tartagal; S22°30’58.9’’ W63°48.079’) and Buenos Aires (La Plata; S 34°55’17.2’’ W 57°57.272’; Merlo (34°39′55″S 58°43′39″O); Arturo Seguí (34°53′16″S 58°07′36″O); Lomas de Zamora (34°46′00″S 58°24′00″O); Avellaneda (34°40′00″S 58°21′00″O); Quilmes 34°43′00″S 58°16′00″O; La Matanza 34°43′00″S 58°38′00″O and Tigre (34°25′00″S 58°35′00″O) (see map in Figure 1). The tested mosquitoes were captured as immature stages or adults from the field. Immature stages were collected from artificial containers. All stages were reared to adults in the laboratory facilities of Centro de Estudios Parasitológicos y de Vectores (CEPAVE). Adults also were captured in the field with aspirators, nets, or traps. Morphological identification was determined under a stereoscopic microscope using dichotomous taxonomic keys [26]. Mosquito samples were conserved in ethanol at -80 °C until genomic DNA was extracted from individuals. For DNA extractions an adapted protocol based on magnetic nanoparticles was used [27]. DNA from each individual was eluted in 50 µl water.

**Figure 1:**
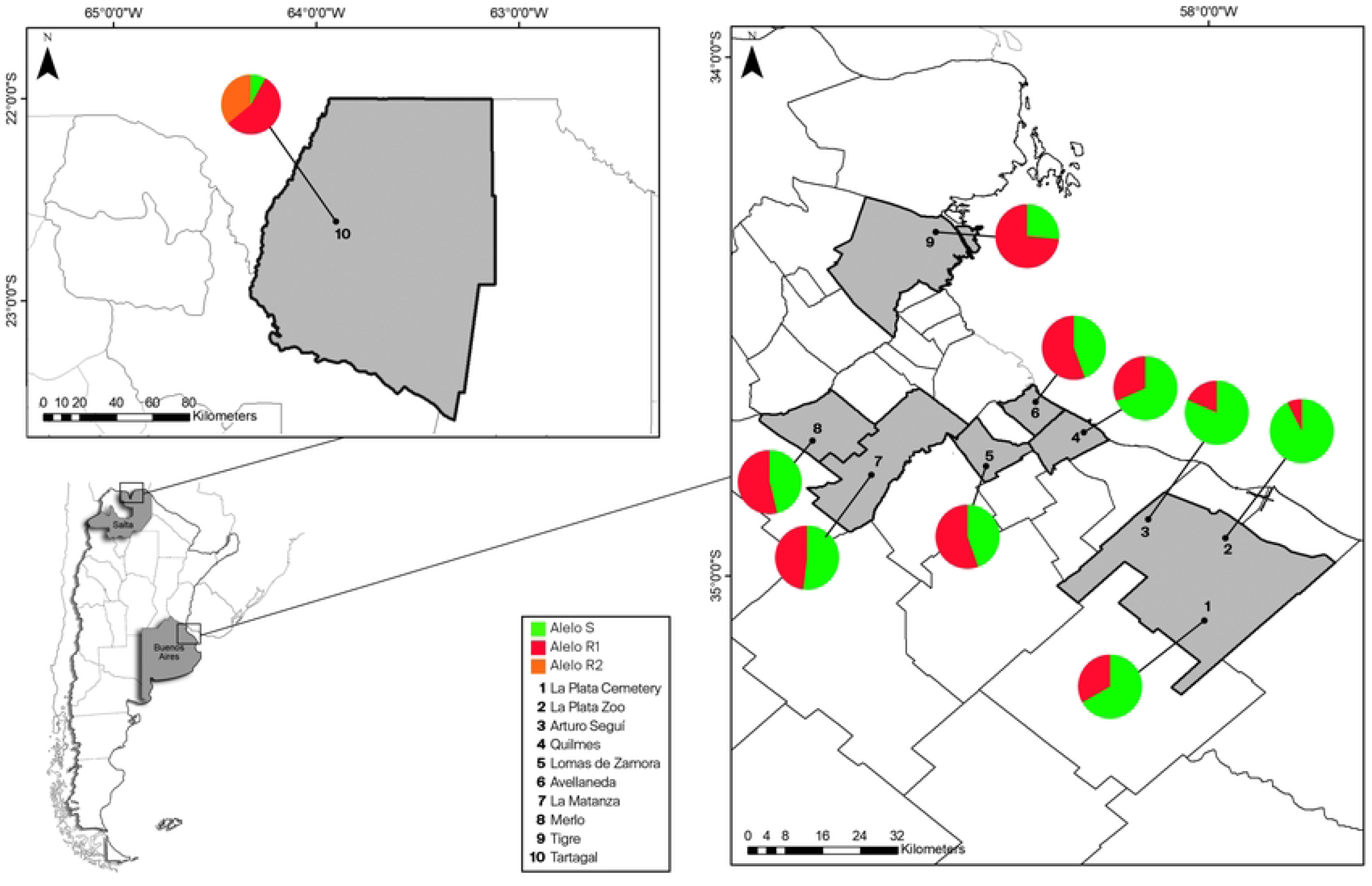
Distribution of the kdr genotypes in Tartagal (Salta Province; upper insert) and Metropolitan Buenos Aires region (MABA; right insert).

### High resolution melting development

The Primers3 v.0.4.0 software (http://bioinfo.ut.ee/primer3-0.4.0/primer3/) was used for the primer design flanking the IIS6 and IIIS6 Na_V_ segments, which contain the positions 1016 and 1534, respectively, in the *aedaena*_*v*_ gene (see primer sequences in Table 1). Amplicons’ melting temperatures (MT) were predicted with the uMELT online software (https://www.dna.utah.edu/umelt/umelt.html). GC tails were added to forward and reverse primers of 1534 amplicon in order to differentiate in at least 2 °C MT of both amplicons. Both the RT-PCR and HRM steps were performed on the ArialMx Real time PCR instrument (Agilent). PCR was performed in 12 μL containing 6 μL Kapa HRM-Fast Master mix (Roche), supplemented with 1.5 mM MgCl_2_, 200 nM of each primer and 2 μL of genomic DNA diluted 1/10. The PCR amplification protocol began with a denaturation step at 95 °C for 3 minutes, followed by 40 cycles of 5 seconds denaturation at 95 °C and 30 seconds annealing-extension at 62 °C. After amplification, the reaction tubes were cooled-down to 55 °C; then warmed from 55 °C to 95 °C at the rate of 0.2 °C/second. The melting curves were analyzed with AriaMX 1.5 software (Agilent). For the setting of the reaction, samples of known genotype: Na_V_SS (1016 VV +1534 FF), Na_V_R2S (1016 VI + 1534 FC) and Na_V_R2R2 (1016 II + 1534 CC)) were used. Samples of known genotype were also included in every mHRM plate for comparisons with samples of unknown genotype.

**Table 1:**
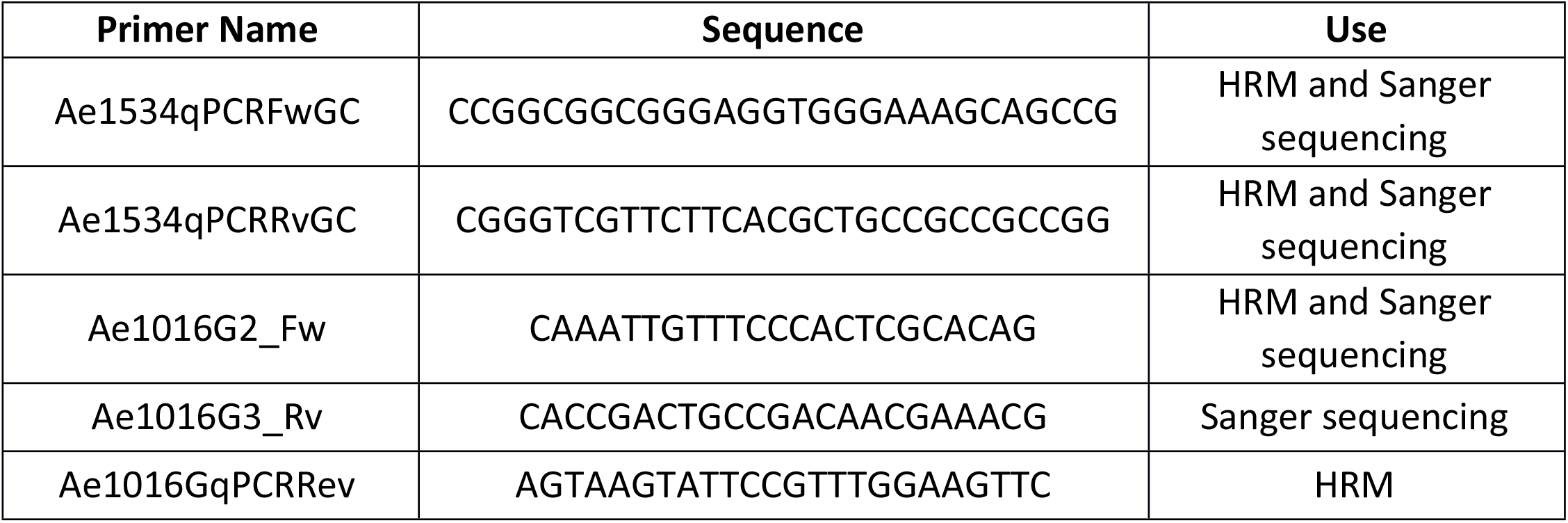

### Sanger sequencing

Genotypes of random selected samples were confirmed by sequencing of the relevant regions of the *aedaenav* gene. For Sanger sequencing, the genomic DNA was used as a template for the PCR reactions using Go-Taq (Promega). The cycling consisted in initial denaturation: 5 min at 95°C; 40 cycles: 30 s at 95°C (denaturation), 30 s at 62°C, and a final extension cycle at 72°C for 5 min. Samples were sequenced for 1016 amplicon and 31 for 1534 amplicon by Sanger method at Macrogen (Seoul-Korea), using Ae1016G3_rev and Ae1534qPCR_Fw primers respectively (Table 1).

## Results

Fragments containing the 1016 and 1534 positions of the *aedaenav* gene were simultaneously amplified in duplex PCR from genomic DNA, using two primer pairs in the reaction tube. By the addition of GCs tails in 5’ regions of forward and reverse primers flanking 1534 position (Table 1), we achieved a difference of around 12 °C in the MT between 1016 (MT around 73.5°C) and 1534 (MT around 85.5°C) amplicons (Figure 2A). This difference allowed the individual analysis of both polymorphic sites using two primer pairs in a single reaction. As expected for a substitution of a guanine (in the wild type V1016) by an adenine (in the 1016I^kdr^ variant), the MTs of the IIS6 amplicons showed its higher value for the 1016 VV genotype (74.23 °C +/-0.03 °C; N=6) and the lower for the 1016 II (73.11 °C +/-0.02 °C; N=8), whereas the heterozygote 1016 VI presented an intermediate MT value (73.88 °C +/-0.04 °C; N=12) (Figure 2 B and C). On the other hand, in the IIIS6 segment MTs were 85.47 °C (+/-0.06°C, N=6) for 1534 FF and 85.93 °C (+/-0.02°C, N=7) for 1534 CC. The heterozygote 1534 FC presented an intermediate MT value (85.64°C +/-0.03°C, N=14) (Figure 2D and E).

**Figure 2:**
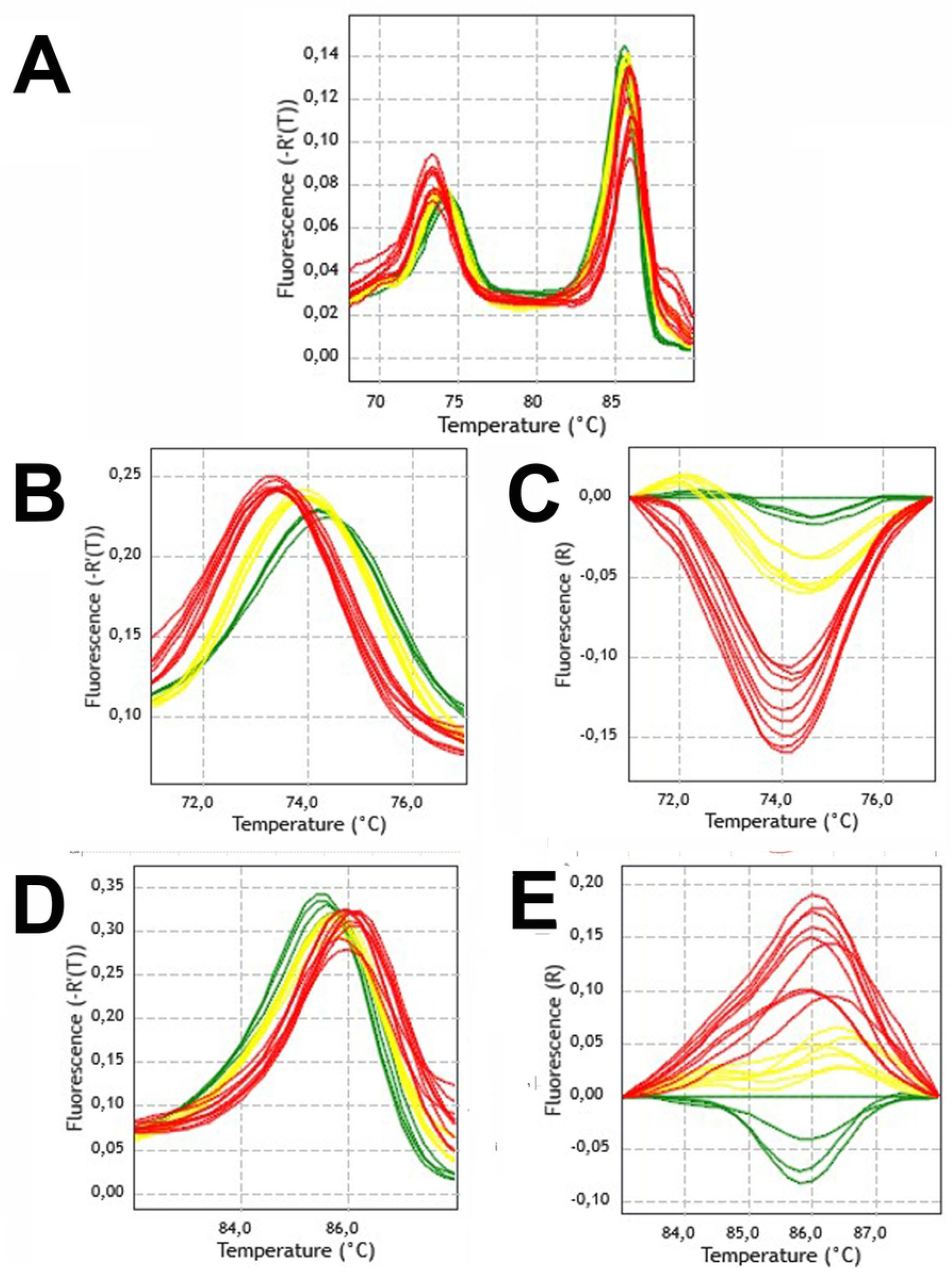
Results of genotyping 1016 and 1534 position in *aedaenav* gene with HRM. Detection of kdr single nucleotide polymorphism by melt curve analysis. Alleles are distinguished by the changes in the melting temperature. **A:** Raw results of mHRM. Peaks on the left and on the right belong respectively to IIS6 and IIIS6 Na_V_ segment amplicons. **B-C:** Derivative and difference melting plots for IIS6 with variation in the 1016 position. **D-E:** Derivative and difference melting plots for IIIS6 with variation in the 1534 position. Red: *kdr* homozygous standard; Yellow: heterozygous standard; Green: wild-type homozygous standard.

We obtained MTs for the IIS6 and IIIS6 paired amplicons with their respective variations in 1016 and 1534 positions, for the six possible genotypes (Table 2; Figure 3). Discernible MTs were observed for genotyping of homozygous in both positions. As expected, the genotyping of heterozygous samples, particularly in position 1534, could be less straightforward for particular samples. We observed that quality and quantity of DNA extracted from individual mosquitoes is a key factor for a correct genotyping, especially for the heterozygous samples.

**Table 2:** MTs ± SEM for each genotype in positions 1016 and 1534.

| Table 2: MTs ± SEM for each genotype in positions 1016 and 1534 | | | | |
| --- | --- | --- | --- | --- |
| genotype | | | Melting temperature ($^{\circ}$C) ± SEM | |
| | 1016 | 1534 | 1016 ($\Delta$ to SS) | 1534 ($\Delta$ to SS) |
| SS | VV | FF | 74.23 ± 0.03 (0.00) | 85.47 ± 0.06 (0.00) |
| SR1 | VV | FC | 74.16 ± 0.02 (0.07) | 85.49 ± 0.02 (0.02) |
| SR2 | VI | FC | 73.88 ± 0.04 (0.35) | 85.64 ± 0.03 (-0.17) |
| R1R1 | VV | CC | 74.16 ± 0.04 (0.07) | 85.86 ± 0.08 (-0.39) |
| R1R2 | VI | CC | 73.45 ± 0.05 (0.78) | 85.94 ± 0.03 (-0.47) |
| R2R2 | II | CC | 73.11 ± 0.02 (1.12) | 85.93 ± 0.02 (-0.46) |

**Figure 3:**
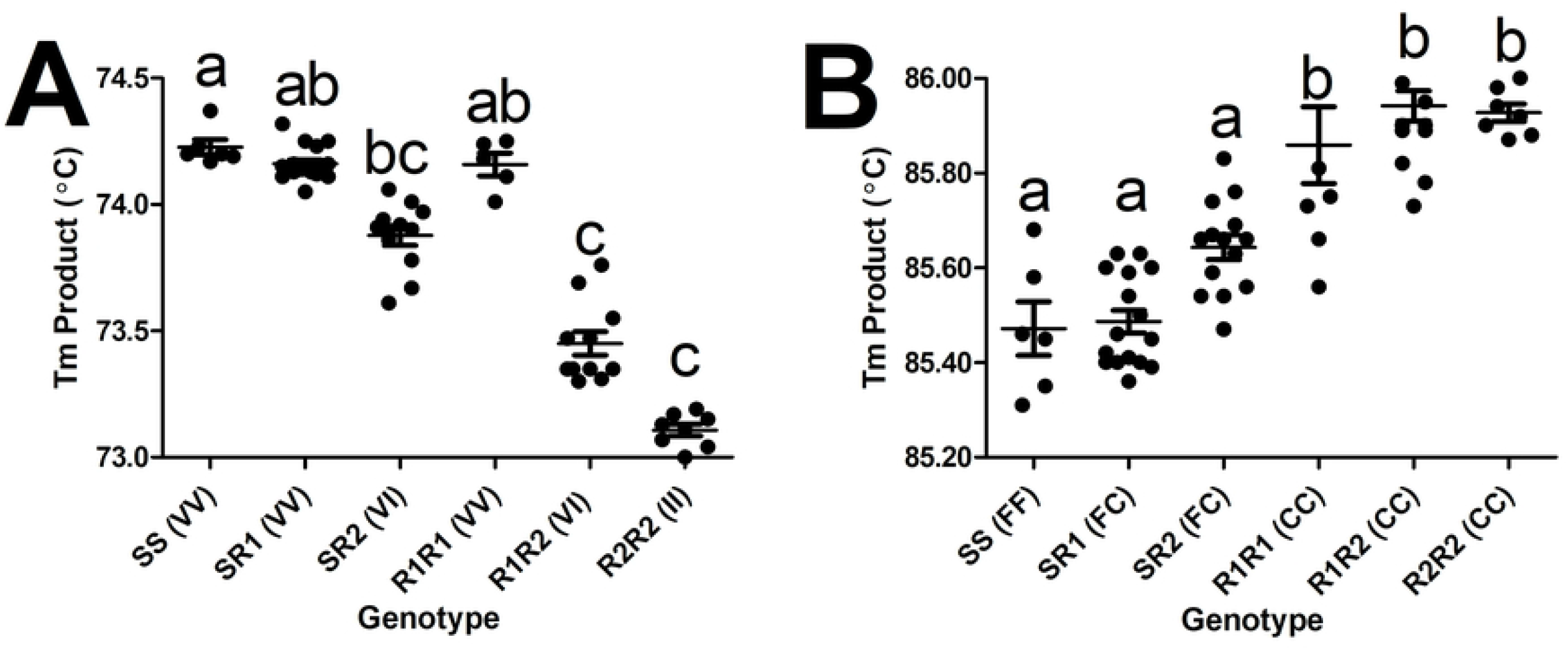
mHRM TMs for the six possible genotypes, considering the variations in the 1016 (A) and 1534 (B) Na_V_ positions in *Aedes aegypti*. Columns marked with different letters are significantly different (non-parametric ANOVA, Dunn’s Multiple Comparison Test, p<0.05)

Table 3 compares different methods available for kdr genotyping in *Ae. aegypti* [28,29]. We found that the mHRM method developed here presented advantages compared to the other options: higher throughput, one step and closed tube, given that both alleles are detected in a single reaction.

**Table 3.** Comparison of methods for the genotyping of KDR mutations in 1016 and 1534 positions^a^.

|  |  |  |  |  |
| --- | --- | --- | --- | --- |
|  | Multiplex allele-specific PCR<sup>b</sup> | Taqman<sup>c</sup> | Singleplex HRM | Multiplex HRM |
| Equipment required | Thermocycler; electrophoresis system; transilluminator with photo camera; computer | Real-time thermocycler; computer | Real-time thermocycler; computer | Real-time thermocycler; computer; software |
| Equipment cost (USD) | 9,500 | 20,000 | 20,000 | 21,000 |
| Time for completion | 4.5 hours | 2.5 hours | 2.5 hours | 1.2 hours |
| Characteristics | PCR followed by gel electrophoresis; multiplex | closed tube; one independent reaction for each mutation | closed tube; one independent reaction for each mutation | closed tube; multiplex |
| Number of steps | 2 | 2 | 2 | 1 |
| number of primers | 7 | 4 | 4 | 4 |
| number of TaqMan probes | 0 | 4 | 0 | 0 |
| number of tubes/wells required | 96 | 192 | 192 | 96 |
| Running cost (USD) | 40 | 212 | 129 | 64 |
| <sup>a</sup> for genotyping 85 individuals in both positions |  |  |  |  |
| <sup>b</sup> Saingamsook et al 2017. |  |  |  |  |
| <sup>c</sup> Macoris et al 2018 |  |  |  |  |
| <sup>d</sup> costs are estimated considering prices in Argentina |  |  |  |  |

We used mHRM for the analysis of *Ae. aegypti* from eight populations from the Center region of Argentina (Metropolitan Area of Buenos Aires; MABA) and one population in the North-West (Tartagal-Salta Province) (Figure 1). Our results show that R1 and S alleles were present in all the localities under study at different frequencies, whereas R2 was only found in Tartagal (Table 4; Figure 1). In order to confirm the mHRM results by a gold standard sequencing method, a number of samples of genomic DNA were PCR amplified for IIS6 and IIIS6 Na_V_ segments with specific primers and submitted to direct Sanger sequencing. For the IIS6 segment, 100% of the sanger sequenced samples were in agreement with mHRM (N=16). All the sequenced IIS6 fragments were identical, with the exception of samples from Tartagal that presented the *kdr* substitution in the 1016 position (Supplementary information 1_A). For the IIIS6 segment, results of mHRM and sanger were coincident in 21 out of 23 samples, in which the 2 misclassified sequences carried a silent substitution of thymidine to cytosine in position 1528 (Supplementary information 1_B) that affected MT of the amplicon. This polymorphism in the 1528 position was previously reported in samples from Africa and the Americas [13].

**Table 3.** Genotypes of *Aedes aegypti* populations considering the V1016I and F1534C kdr mutations.

|  |  |  |  |  |  |
| --- | --- | --- | --- | --- | --- |
| Region | Province-Locality | Type or population | Number of individuals | Genotype | Frecuence (%) |
| Center | La Plata Zoo-Buenos Aires | Peri-urban | 20 | SS | 85 |
|  |  |  |  | SR1 | 15 |
| Center | La Plata Cemetery-Buenos Aires | Peri-urban | 30 | SS | 43.3 |
|  |  |  |  | SR1 | 46.7 |
|  |  |  |  | R1R1 | 10 |
| Center | Arturo Seguí-Buenos Aires | Rural | 28 | SS | 64.3 |
|  |  |  |  | SR1 | 35.7 |
| Center | Quilmes-Buenos Aires | Urban | 30 | SS | 46.7 |
|  |  |  |  | SR1 | 43.3 |
|  |  |  |  | R1R1 | 10 |
| Center | Lomas de Zamora-Buenos Aires | Urban | 28 | SS | 21.4 |
|  |  |  |  | SR1 | 46.4 |
|  |  |  |  | R1R1 | 32.1 |
| Center | Avellaneda-Buenos Aires | Urban | 27 | SS | 14.8 |
|  |  |  |  | SR1 | 63 |
|  |  |  |  | R1R1 | 22.2 |
| Center | La Matanza-Buenos Aires | Urban | 27 | SS | 18.5 |
|  |  |  |  | SR1 | 66.7 |
|  |  |  |  | R1R1 | 14.8 |
| Center | Merlo-Buenos Aires | Urban | 26 | SS | 30.8 |
|  |  |  |  | SR1 | 30.8 |
|  |  |  |  | R1R1 | 38.5 |
| Center | Tigre-Buenos Aires | Urban | 30 | SS | 16.7 |
|  |  |  |  | SR1 | 20 |
|  |  |  |  | R1R1 | 63.3 |
| North West | Tartagal-Salta | urban | 25 | SR1 | 8 |
|  |  |  |  | SR2 | 8 |
|  |  |  |  | R1R1 | 28 |
|  |  |  |  | R1R2 | 48 |
|  |  |  |  | R2R2 | 8 |

In MABA, we analyzed both rural/peri-urban locations and populated urban localities. In the former, the majoritarian allele was S (92.5% in La Plata Zoo; 81.5% in Arturo Seguí and 66.7% in La Plata Cemetery). In the urban localities, a higher frequency of R1 was detected, being minority in Quilmes (31.7%), but majority in Tigre (73.4%). In the other localities analyzed, the rate of R1 allele was around 50.0% (55.4% in Lomas de Zamora, 55.6% in Avellaneda, 48.1% in La Matanza and 53.85% in Merlo) (Figure 1; Table 3). In Tartagal, a locality with a more intense historical use of pyrethroids, R2 allele was detected in 36.0% of the samples, despite the majoritarian allele was R1 (56.0%), and the S allele was found in only 8.0% of the individuals.

## Discussion

The development of pyrethroid resistance in *Ae. aegypti* seriously compromises dengue control campaigns. Alternative adulticides like the organophosphate Malathion are not admitted for treatment of human dwellings in Argentina, given environmental and sanitary considerations [8]. The inclusion of resistance-management strategies is necessary in rational vector control campaigns, in order to prolong the effective life of pyrethroids. In this context, the monitoring of resistance-conferring alleles, such as kdr mutations, should be performed as a routine practice to aid in the decision-making process on control campaigns. Notwithstanding the wide distribution of kdr alleles in *Ae. aegypti* natural populations all over the world, and the epidemiologic situation in Argentina, kdr presence and/or distribution was not previously reported in this country.

There are a number of assays available for genotyping kdr alleles in *Ae. aegypti*. The most widely used of these for populations in the American continent is based on Taqman probes [15,29]. Allele-specific PCR was also developed; even though lower in cost, this technique has also a lower throughput, and the reaction conditions could not be reproducible in different labs, and must be carefully determined in order to avoid incorrect results [23]. Singleplex HRM was used for genotyping the mutations present in Asian populations of *Ae. aegypti* [28], doubling the genotyping efforts when compared with the mHRM proposed here. Multiplex allele-specific PCR could be a suitable option for lower equipped laboratories, even though skilled labor costs should be considered. Mosquitos in all the life stages (from eggs to adults) can be sampled during routine surveillance activities, even dead individuals, without requiring efforts to collect or rearing insects. Using a high-throughput genotyping technique such as the mHRM presented here, the results could be obtained in a few hours (Table 2). Careful in sample conservation and the use of good yield DNA extraction methods are strongly recommended. In cases of ambiguous genotyping results, mainly for heterozygous samples, the use of a complementary method (such as TaqMan probes or Sanger sequencing) can be implemented. In this way, the complemented use of available methods, including the mHRM presented here, will improve genotyping efforts in terms of time and economic costs.

Despite there is no detailed information about the insecticide application by region in Argentina, we can assume a direct relationship between dengue epidemics and the adult control using spatial sprays with adulticides, as is recommended during the dengue outbreaks by the National Health Ministry. Interestingly, in Tartagal (Salta), a region under a prolonged use of pyrethroids, since dengue epidemics in Salta Province are recorded from 1998 [30] with reported cases almost every year [31] until present (Bulletin from the Ministry of Health Argentina, 2022), we observed a higher frequency of R2 genotype. In agreement, Harburger et al [32] reported for the first time pyrethroid resistance in *Ae. aegypti* adults from Argentina, in Salta Province (Salvador Mazza). Buenos Aires Metropolitan Area has a more recent history of dengue outbreaks, in despite of being the most populated region from Argentina, representing 35% of the total population with two important dengue outbreaks registered in 2016 [33] and 2020 (Bulletin from the Ministry of Health Argentina. 2020). Interestingly, a positive correlation exists between the years of treatment with pyrethroids and the emergence of V1016I^kdr^ mutation. Also, those rural or peri-urban populations, not treated with pyrethroids in their domi-sanitary use, presented a higher proportion of the wild-type (sensitive) allele. These results are in agreement with a sequential selection of kdr mutations in *Ae. aegypti* [19]. Given that V1016I^kdr^ mutation alone does not confer insecticide resistance, mutation F1534C^kdr^ seems to emerge earlier in response to the selective pressure with pyrethroids. V1016I^kd^+ F1534C^kdr^ allele emerged in the Americas more recently, leading to a greater and broader spectrum of pyrethroid resistance.

Even though the frequency results obtained here should be interpreted cautiously, as a greater number of individuals would be analyzed, they indicate for the first time the extended presence of *kdr* alleles in field populations from Argentina. Also, we show that these alleles are present in distant regions, suggesting a wide distribution. The information on *kdr* alleles in Buenos Aires and Salta Provinces should be considered for rational decisions in vector control campaigns in Argentina. Particularly in the populated MABA region, present results should be taken as a warning signal in the design of *Ae. aegypti* control campaigns. The treatments with pyrethroids should be carefully and rationally planned, in order to extend the utility of these compounds for longer periods.

## Conclusions

We have developed a high-throughput multiplex HRM method for the genotyping of alleles associated with pyrethroid resistance in *Ae. aegypti* from the American continent. The method developed here is comparable in its sensitivity and reliability with other genotyping methods, but reduces costs and running time. It could be incorporated in control campaigns for control the presence and spreading of resistance-associated alleles, in order to reduce costs and running times; optimal results will be obtained with the complement of several available genotyping methods. We report here for the first time the presence of kdr mutations in distant populations from Argentina, with different epidemiological situations and different history of mosquito control efforts.

## Acknowledgements

Authors would like to thanks to Claudia Rodriquez Torres, Pedro Mendoza Zélis, Elisa de Sousa and Luciana Juncal for their generous providing of silica-coated magnetic nanoparticles for DNA extractions, Laura Morote for helping in designing the figures, Darío Balcazar from the molecular laboratory at CEPAVE and Melisa B. Bonica, who collected mosquito samples in the context of hers PhD Thesis. The Argentinean groups are members of Argentinean Network for the Research in Pesticide Resistance (RAREP).

